# Genome-wide reversal of glial and neurovascular cell transcriptomic aging signatures by GLP-1R agonism

**DOI:** 10.1101/2020.12.21.423879

**Authors:** Zhongqi Li, Xinyi Chen, Joaquim S. L. Vong, Lei Zhao, Junzhe Huang, Leo Y. C. Yan, Bonaventure Ip, Yun Kwok Wing, Hei-Ming Lai, Vincent C. T. Mok, Ho Ko

**Author notes:** **Address correspondence to:** Vincent C. T. Mok, Department of Medicine and Therapeutics, Faculty of Medicine, CUHK, Shatin, Hong Kong.; *or* Ho Ko, Department of Medicine and Therapeutics, Faculty of Medicine, CUHK, Shatin, Hong Kong. These authors contributed equally to this work (Zhongqi Li, Xinyi Chen, Joaquim S. L. Vong, Lei Zhao).

## Abstract

Pharmacological reversal of brain aging is a long-sought yet challenging strategy for the prevention and treatment of age-related neurodegeneration, due to the diverse cell types and complex cellular pathways impacted by the aging process. Here, we report the genome-wide reversal of transcriptomic aging signatures in multiple major brain cell types, including glial and mural cells, by glucagon-like peptide-1 receptor (GLP-1R) agonist (GLP-1RA) treatment. The age-related expression changes reversed by GLP-1RA encompass both shared and cell type-specific functional pathways that are implicated in aging and neurodegeneration. Concomitantly, Alzheimer’s disease (AD)-associated transcriptomic signature in microglia that arises from aging is reduced. These results show the feasibility of reversing brain aging by pharmacological means, provide mechanistic insights into the neurological benefits of GLP-1RAs in diabetic patients, and imply that GLP-1R agonism may be a generally applicable pharmacological intervention for patients at risk of age-related neurodegeneration.

## Introduction

Aging has long been considered irreversible. Nearly all cellular processes are implicated in or impacted by aging, ranging from metabolism, stress response, immune responses, cellular senescence, to gene expression and genomic stability(1). These complex molecular changes presumably lead to an alteration of cellular states and compositions in body organs(1, 2), manifesting as age-related functional decline. Given the complexity of biological changes involved and a lack of easily targetable sets of driving pathways, anti-aging pharmacotherapy is considered highly challenging, if feasible at all. However, it remains an attractive pursuit for tackling age-related disorders, such as neurodegeneration for which aging is the strongest risk factor. Slowing down or even reversing transcriptomic and functional alterations in the aging brain may provide a strategy for the primary prevention and even the treatment of neurodegenerative diseases.

Glucagon-like peptide-1 (GLP-1) is a peptide hormone produced peripherally by the intestinal L-cells for potentiating glucose-dependent insulin release, and centrally in the brain by preproglucagon neurons in the nucleus tractus solitarii(3). In the past decade, multiple pharmacokinetically optimized GLP-1 receptor (GLP-1R) agonists (GLP-1RAs) have been approved for the clinical treatment of diabetes mellitus. Remarkably, recent clinical studies provided compelling evidence that GLP-1RAs exhibit neuroprotective effects beyond that conferred by glycemic control, reducing the incidences of cognitive decline and Parkinson’s disease (PD) in diabetic patients(4, 5). Additionally, GLP-1RAs may slow the progression of established Alzheimer’s disease (AD) and PD in non-diabetic patients(6, 7). Mechanistically, apart from alleviating neuroinflammation in animal models of neurodegeneration(8, 9), we recently demonstrated that exenatide (a GLP-1RA) treatment partially reverses age-related transcriptomic changes in brain endothelial cells (ECs) and reduces nonspecific blood-brain barrier (BBB) leakage(10). How GLP-1RA treatment impacts glial and other neurovascular cell types, whose age-related expression changes also play crucial roles in brain aging and degeneration, remained unclear.

## Results and Discussion

We performed single-cell transcriptomic profiling experiments in young adult, aged and exenatide-treated aged mice (**Fig. 1a, b, Supplementary Fig. 1**), to examine the genome-wide expression changes in glial and neurovascular cells in aging and their modulation by GLP-1R agonism. We calculated significant differentially expressed genes (DEGs, defined as false discovery rate (FDR)-adjusted *P*-value < 0.05) for each cell type, to analyse their patterns of expression changes in aging and the effects of exenatide treatment. In line with our previous report(10), the aging-associated transcriptomic changes in brain ECs were partially reversed by exenatide treatment (**Fig. 1c, d**). Strikingly, the transcriptomic reversal effect of the GLP-1RA was even more profound in other brain cell types (**Fig. 1c, d**). These included multiple glial (**Fig. 1c, d**, AC: astrocyte; OPC: oligodendrocyte precursor cell; MG: microglia) and mural (**Fig. 1c, d**, SMC: smooth muscle cell; PC: pericyte) cell types. This phenomenon was also observed, although weaker, in oligodendrocyte (OLG) and perivascular macrophage (MAC) (**Fig. 1c, d**). Overall, the reversal effect on glial cell transcriptomic aging signatures appeared to be the most prominent in AC and OPC, followed by MG (**Fig. 1d**). Among vascular cells, the effects were stronger in SMC and PC compared with that in EC (**Fig. 1d**).

**Fig. 1.**
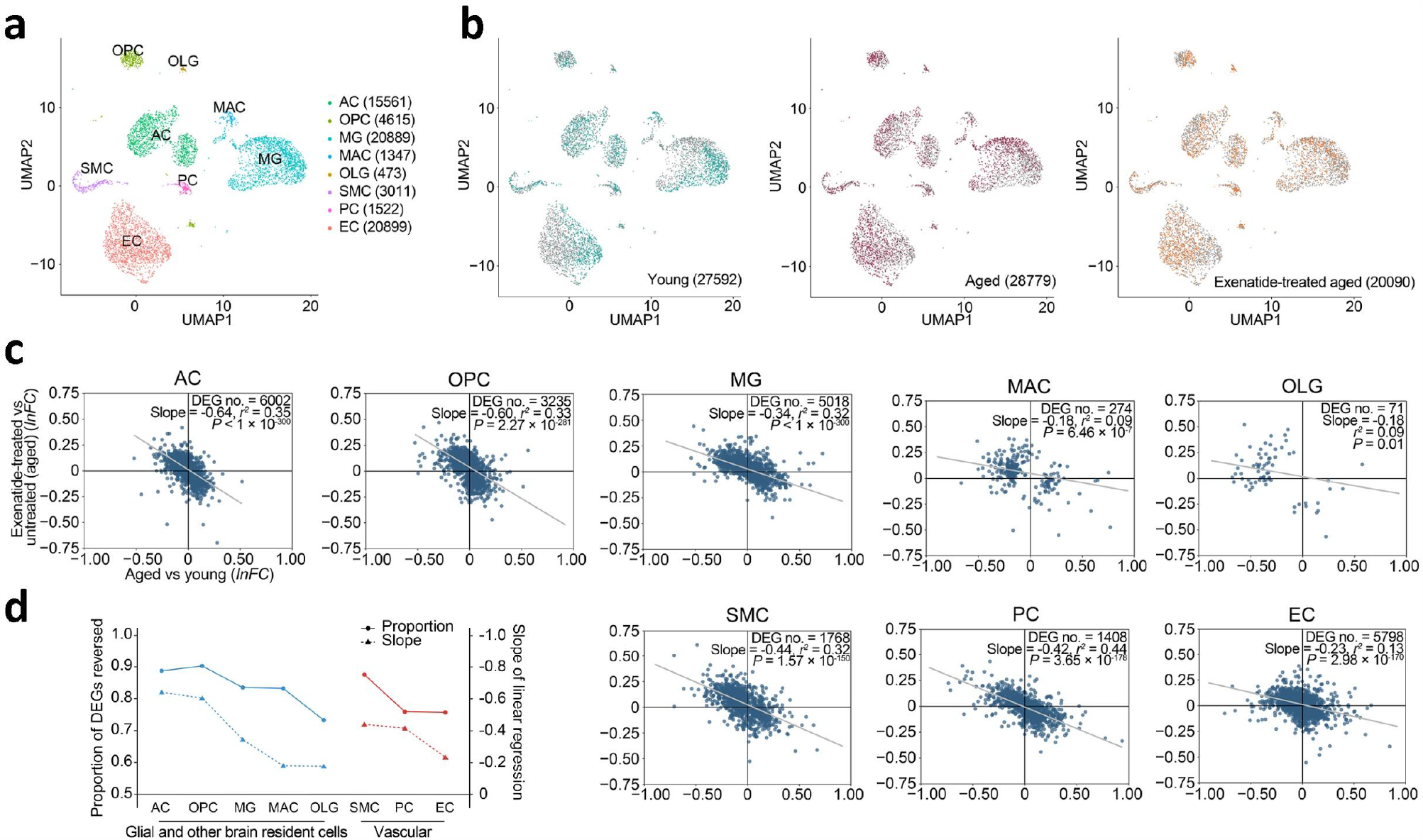
Reversal of glial and neurovascular cell transcriptomic aging signatures by GLP-1RA treatment. **(a)** UMAP visualization of the major cell type clusters identified and analyzed. Numbers in brackets: cell numbers for the respective cell types. Abbreviations: AC: astrocyte; OPC: oligodendrocyte precursor cell; MG: microglia; MAC: perivascular macrophage; OLG: oligodendrocyte; SMC: smooth muscle cell; PC: pericyte; EC: endothelial cell. **(b)** UMAP visualization of the single-cell transcriptomes from young adult, aged and GLP-1RA (exenatide)-treated aged mouse brains. For each plot, colored dots highlight cells from the respective labelled group, while grey dots are cells from the other groups. Numbers in brackets: cell numbers for the respective groups (*n* = 3 animals for each group). For clarity, 6000 cells were subsampled for visualization in each plot in **(a)** and **(b). (c)** Age-related expression changes (*x*-axis) plotted against post-GLP-1RA treatment expression changes (*y*-axis) in glial (AC, OPC, MG and OLG), vascular (EC, PC, SMC) cell types, and MAC. Each dot represents one DEG. *lnFC*: natural log of fold change. Grey lines: lines of best fit by linear regression. **(d)** Proportions of DEGs reversed and the slopes of lines of best fit by linear regression shown in **(c)** in the different cell types.

We next asked if the expression changes reversed by GLP-1RA treatment may be functionally relevant. Pathway enrichment analysis for each cell type on their most prominent reversed DEGs highlighted the amelioration of age-related expression changes involved in extensive cellular functions (**Fig. 2a**). In most cell types, these included genes mediating glucose/energy, lipid and protein metabolic processes (**Fig. 2a**), as well as transcriptional and translational regulation (**Fig. 2a**). We also noted cell type-specific changes by pathway analysis and examining the expression changes of selected genes with significant functional roles. In AC, age-related expression changes in genes mediating immune response, homeostatic functions and synaptic plasticity occur(11). In our dataset, the ACs from the exenatide-treated aged mice appeared to partially revert to a younger phenotype, with downregulation of several complement component 1q genes (**Fig. 2b**), and upregulation of subsets of genes encoding synaptic modification-related proteins, metabolite receptors and transporters, neurotransmitter receptors and ion channels (**Fig. 2b**). On the other hand, immune response and cytokine signaling-related genes were especially prominent among DEGs reversed in MG and MAC, followed by EC (**Fig. 2a**). Indeed, previous studies reported that MGs in the aging and AD brain exhibit pro-inflammatory phenotypes(1, 12–14), with a subset of MGs in AD characterized by a disease-associated microglia (DAM) transcriptomic state(1, 14). After exenatide treatment, the MGs showed an upregulation of multiple homeostatic function-related and immune response inhibitory genes (**Fig. 2b**), reversed expression changes of several activation-associated genes (**Fig. 2b**), and decreased AD-associated MG signature(1, 14) scores (**Fig. 2c, d**). In SMC, we found post-treatment reversal of expression changes involving important functional processes associated with age-related vascular stiffening, such as calcium signaling, extracellular matrix (ECM) remodelling and contractile pathways (**Fig. 2a, b**).

**Fig. 2.**
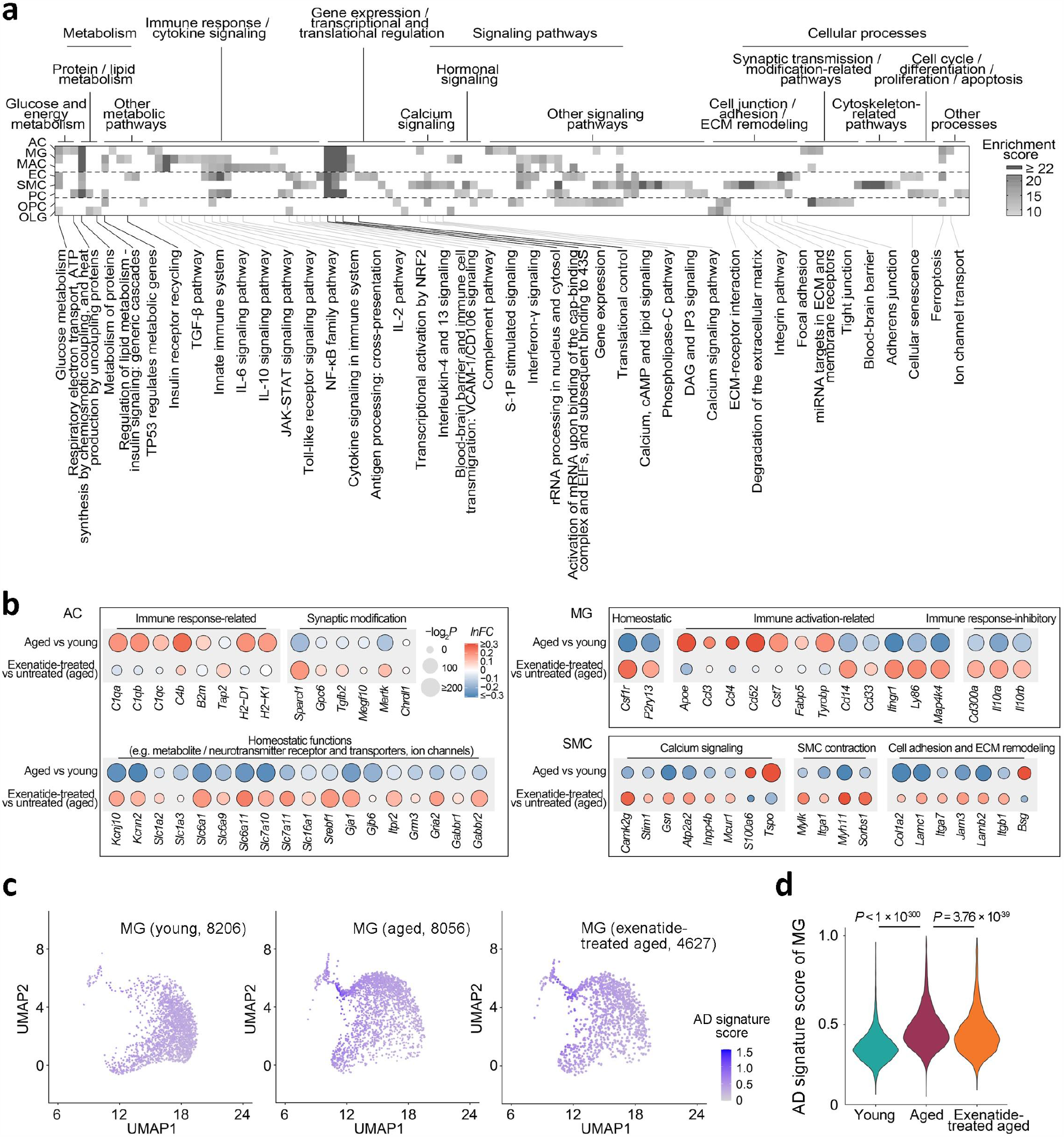
Functional pathway analysis of the GLP-1RA treatment-reversed age-related transcriptomic changes. **(a)** Pathways with significant enrichment among the most prominent age-related expression changes reversed by GLP-1RA treatment in the different cell types. Cell type abbreviations: same as in **Fig. 1. (b)** Dot plots of expression changes of selected functionally important genes in AC, MG and SMC, in aging and post-GLP-1RA treatment. **(c)** UMAP visualization and **(d)** violin plots of AD signature scores in the MGs from young adult, aged and GLP-1RA-treated aged mouse brains. Comparison of AD signature scores in the MGs among the three groups: *P* = 2.2 × 10^−16^, Kruskal-Wallis test; the *P*-values of pairwise comparisons by post hoc Dunn’s test are shown in **(d)**.

As we found the most prominent transcriptomic reversal effect in AC (**Fig. 1c, d**) and AC exhibits molecular phenotypic differences across brain regions(15), we asked if the age-related transcriptomic alterations and the GLP-1RA treatment effect in AC may exhibit regional specificity. We identified four main regional AC subtype clusters, namely telencephalic AC clusters 1 and 2 (ACTE1 and ACTE2 respectively), and non-telencephalic AC clusters 1 and 2 (ACNT1 and ACNT2 respectively), using the expression patterns of known regional AC marker genes(15) (**Supplementary Fig. 2a, b**). Despite differences in their molecular phenotype (**Supplementary Fig. 2a, b**), the AC subtypes shared highly consistent age-related differential expressions (**Supplementary Fig. 3a**). The expression changes induced by exenatide treatment were generally even more consistent in the AC subtypes (**Supplementary Fig. 3a**). Consequently, the transcriptomic reversal effect of exenatide on the aging-associated expression changes was well conserved across all four regional AC subtypes examined (**Supplementary Fig. 3b**). We also performed similar analysis on the mural cell subtypes that segregate along the arteriovenous axis(16) (**Supplementary Fig. 4a, b**). Pairwise comparison of the differential expressions likewise revealed shared age-related expression changes and reversal by GLP-1RA treatment in all four mural cell subtypes (**Supplementary Fig. 5a, b**), despite their intrinsic transcriptomic differences (**Supplementary Fig. 4a, b**).

In conclusion, we demonstrated that a generalized reversal of functionally relevant transcriptomic changes at the genome-wide level in multiple glial and vascular cell types in the aged brain is pharmacologically achievable with GLP-1R agonism. We speculate that the profound pleiotropic effects of GLP-1RA on brain aging may depend on a combination of central and peripheral mechanisms. Centrally in the brain, subsets of GLP-1R-expressing neurons and glial cells may be directly targeted by GLP-1RA(9, 17). Peripherally, GLP-1RA-mediated improvement in metabolic profiles or immunomodulation may impact the glial and neurovascular cells, as they are responsive to compositions of the circulation(18) and interact with peripheral immune cells(19). Clarifying these possibilities will require systematic studies involving knockout or knockdown of GLP-1R in the respective putative target cell types (e.g. neuron, MG). The knowledge will also instruct further developments of small-molecule GLP-1RAs(20–22), which may need to cross the BBB to attain any centrally mediated anti-aging effects in the brain.

## Methods

### Animal subjects

All experimental procedures were approved in advance by the Animal Research Ethical Committee of CUHK and were carried out in accordance with the Guide for the Care and Use of Laboratory Animals. C57BL/6J mice were provided by the Laboratory Animal Service Center of CUHK and maintained at controlled temperature (22 – 23°C) with an alternating 12-hour light/dark cycle with free access to standard mouse diet and water. Male mice of two age groups (2 – 3 months old and 18 – 20 months old) were used for experiments. For the treatment groups, exenatide (5 nmol/kg bw, Byetta, AstraZeneca LP) was intraperitoneally (I.P.) administered daily starting at 17 – 18 months old for 4 – 5 weeks prior to experimentation(10).

### Single-cell dissociation and RNA sequencing

Identical brain tissue digestion, cell dissociation and single-cell RNA sequencing protocols were adopted as per our previous study(10), except the use of Chromium Single Cell 3′ Reagent Kits v3 (10X Genomics, US). Briefly, mice were deeply anaesthetized and perfused transcardially with 20 ml of ice-cold phosphate buffered saline (PBS). Mice were then rapidly decapitated, and whole brains (except cerebellum) were immersed in ice-cold Dulbecco’s modified Eagle’s medium (DMEM, Thermo Fisher Scientific, US). The brain tissues were cut into small pieces and dissociated into single cells using a modified version of the Neural Tissue Dissociation Kit (P) (130-092-628, Miltenyi Biotec, US). Myelin debris was removed using the Myelin Removal kit II (130-096-733, Miltenyi Biotec, US). Cell clumps were removed by serial filtration through pre-wetted 70-μm (#352350, Falcon, US) and 40-μm (#352340, Falcon, US) nylon cell strainers. Centrifugation was performed at 300 ×g for 5 minutes at 4°C. The final cell pellets were resuspended in 500 – 1000 μl FACS buffer (DMEM without phenol red (Thermo Fisher Scientific, US), supplemented with 2% fetal bovine serum (Thermo Fisher Scientific, US)).

Single-cell RNA sequencing libraries were generated using the Chromium Single Cell 3′ Reagent Kit v3 (10X Genomics, US). Briefly, single-cell suspension at a density of 500 – 1,000 cells/μL in FACS buffer was added to real-time polymerase chain reaction (RT-PCR) master mix and then loaded together with Single Cell 3′ gel beads and partitioning oil into a Single Cell 3′ Chip (10X Genomics, US). RNA transcripts from single cells were uniquely barcoded and reverse-transcribed within droplets. cDNA molecules were preamplified and pooled, followed by library construction. All libraries were quantified by Qubit and RT-PCR on a LightCycler 96 System (Roche Life Science, Germany). The size profiles of the pre-amplified cDNA and sequencing libraries were examined by the Agilent High Sensitivity D5000 and High Sensitivity D1000 ScreenTape Systems (Agilent, US), respectively. All single-cell libraries were sequenced with a customized paired-end with single indexing (26/8/98-bp for v3 libraries) format. All single-cell libraries were sequenced on a NextSeq 500 system (Illumina, US) using the NextSeq 500 High Output v2 Kit (Illumina, US). The library sequencing saturation was on average 68.73%. The data were aligned in Cell Ranger (v3.0.0, 10X Genomics, US). Compared to the dataset in our previous study(10), the current one has improved mRNA capture efficiency (median genes per cell: 2,091, range: 505 to 5,791), allowing more sensitive detection of differentially expressed genes (DEGs).

### Data quality control, single-cell transcriptome clustering and visualization

Data processing and visualization were performed using the Seurat package (v3.1.5)(23) and custom scripts in R (v3.6.1). The raw count matrix was generated by default parameters (with the mm10 reference genome). There were 106,902 cells in the primary count matrix. Genes expressed by fewer than five cells were removed, leaving 21,259 genes in total. Among these genes, 3,000 high-variance genes were identified by the Seurat *FindVariableFeatures* function. The dataset was filtered to exclude low-quality cells by the following criteria: (1) < 5% or > 95% UMI count or gene count, or (2) proportion of mitochondrial genes > 20%. Gene count normalization and high-variance gene identification were applied to the raw data of 78,490 cells retained for further analysis. The *SCTransform* function in the Seurat package was used for expression normalization by fitting the data to a negative binomial regression model. For dimensionality reduction, principal component analysis (PCA) was applied to compute the first 30 top principal components. Clustering was carried out by the Seurat functions *FindNeighbors* and *FindClusters*. Modularity optimization was then performed on the shared nearest neighbor graph results from *FindNeighbors* for clustering (resolution parameter: 1.4). We employed Uniform Manifold Approximation and Projection (UMAP)(24) for visualization of the single-cell transcriptomes and clustering results.

### Cell type and subtype identification

To identify primary cell types, we employed known cell type-specific marker genes and examined their expression levels among all initial clusters included. We excluded clusters with dual-high expression of two or more cell type-specific marker genes. These included a cluster with high expression of both endothelial cell and pericyte markers, corresponding to contamination of pericytes by endothelial cell fragments(16, 25, 26). The remaining clusters were classified into primary cell types, whereby 8 were analyzed in this study (see **Fig. 1** and **Supplementary Fig. 1**). Telencephalic and non-telencephalic AC subtypes were identified by unsupervised clustering of the AC transcriptomes (after expression normalization among ACs), followed by examining the expression patterns of regionally specific marker genes(15) (see **Supplementary Fig. 2**). To identify SMC subtypes, the *CellAssign* algorithm was employed(27), using previously reported SMC subtype marker genes(16) as prior.

### Differential expression analysis

The Seurat *FindMarkers* function and the MAST package (v1.8.2) were employed for the calculation of differentially expressed genes (DEGs) with associated false discovery rate (FDR)-adjusted *P*-value and magnitude of change expressed in natural log of fold change (*lnFC*) for each cell type or subtype. We define significant DEGs as those with FDR-adjusted *P*-value < 0.05.

### Transcriptomic reversal effect, pathway enrichment and consistency of differential expressions analyses

For each cell type or subtype, the union of significant DEGs in either or both the aged versus young adult group, and the exenatide-treated versus untreated aged group comparisons were included for transcriptomic reversal analysis. Proportions of genes with opposite directionality of changes (i.e. reversed), linear regression analysis with slope of line of best fit, associated *r*^2^ and *P*-values were obtained. For pathway enrichment analysis of the reversed DEGs of a given cell type, DEGs with fold changes ranking within top 500 in both age- and treatment-stratified comparisons and opposite directionality of changes were included. Pathways with significant enrichment were identified using the GeneAnalytics platform(28). To examine the consistency of differential expressions across cell subtypes, for each pair of cell subtypes, the union of their significant DEGs in a given comparison (i.e. age- or treatment-stratified) were included. Linear regression analysis was then carried out with slope of line of best fit, associated *r*^2^ and *P*-values obtained.

### Calculation of Alzheimer’s disease (AD)-associated microglia signature scores

For each single MG transcriptome, the AD-associated MG signature score was calculated using the *AddModuleScore* function in the Seurate package, as the average normalized expression levels of top 100 upregulated genes in AD-associated MGs(14) subtracted by the aggregated expression of control feature sets, similar to previously described(1).

### Data availability

The single-cell RNA sequencing data has been deposited at the Broad Institute Single Cell Portal [https://singlecell.broadinstitute.org/single_cell/study/SCP1182/glp1ra-brain-aging-reversal]. The R code for data analysis is available at [https://github.com/RichardLZQ/NVUB5_code].

## Acknowledgments

We thank Dennis Lo and Rossa Chiu for generous support; Becky Yung, Florence Yau, Anki Miu and Rebecca Chau for administrative support to the project. This work was funded by a Croucher Innovation Award (CIA20CU01) (H.K.); Faculty Innovation Awards (FIA2020/B/01 and FIA2017/B/01) from the Faculty of Medicine, CUHK (H.M.L. and H.K.); the Collaborative Research Fund (C6027-19GF) and the Area of Excellence Scheme (AoE/M-604/16) of the University Grants Committee of Hong Kong (H.K.).

## Author Contributions

Z.L., X.C., J.S.L.V. and L.Z. carried out the in vivo treatment and single-cell RNA sequencing experiments. L.Y.C.Y. assisted the experiments. Z.L. and X.C. analyzed the data. J.H., B.I., Y.K.W. and H.M.L. contributed to data interpretation. Z.L., X.C., J.H., H.M.L., V.C.T.M. and H.K. wrote the paper with input from all authors.

## Figures

**Supplementary Fig. 1.**
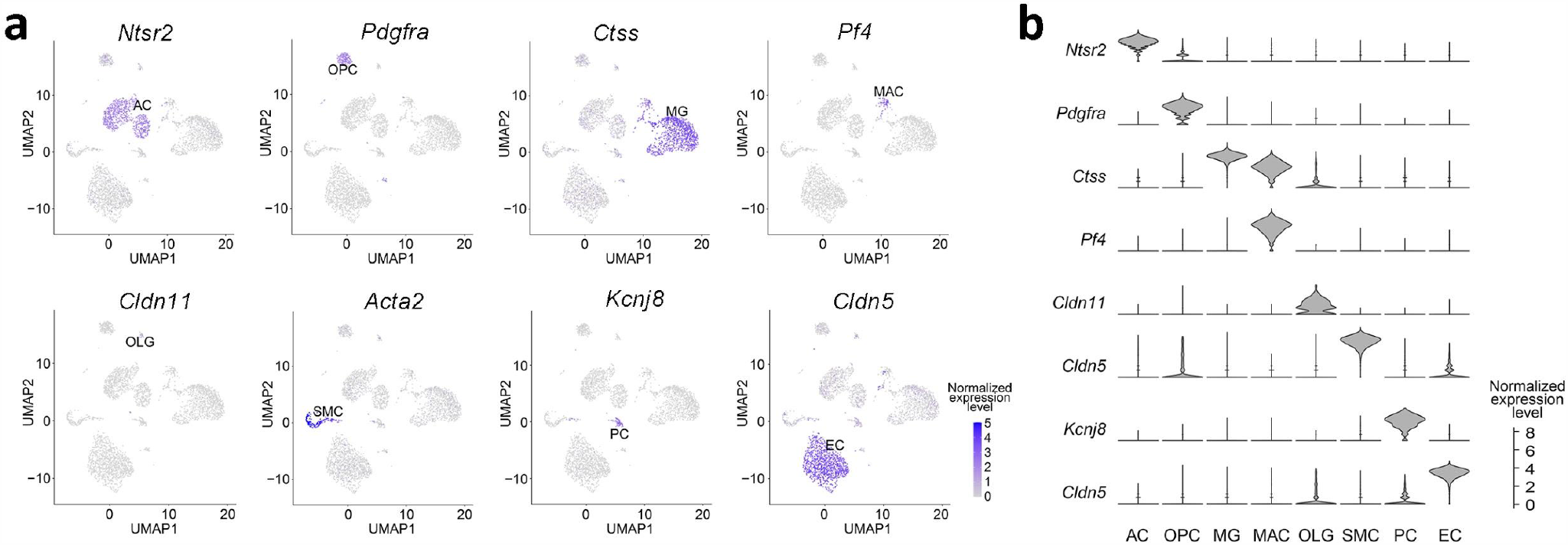
Primary cell type identification and their marker gene expression patterns. **(a)** UMAP visualization and **(b)** violin plots of cell type-specific marker gene expression patterns in the different cell type clusters. Cell type abbreviations: same as in **Fig. 1**. For clarity, 6000 cells were subsampled for visualization in each plot in **(a)**.

**Supplementary Fig. 2.**
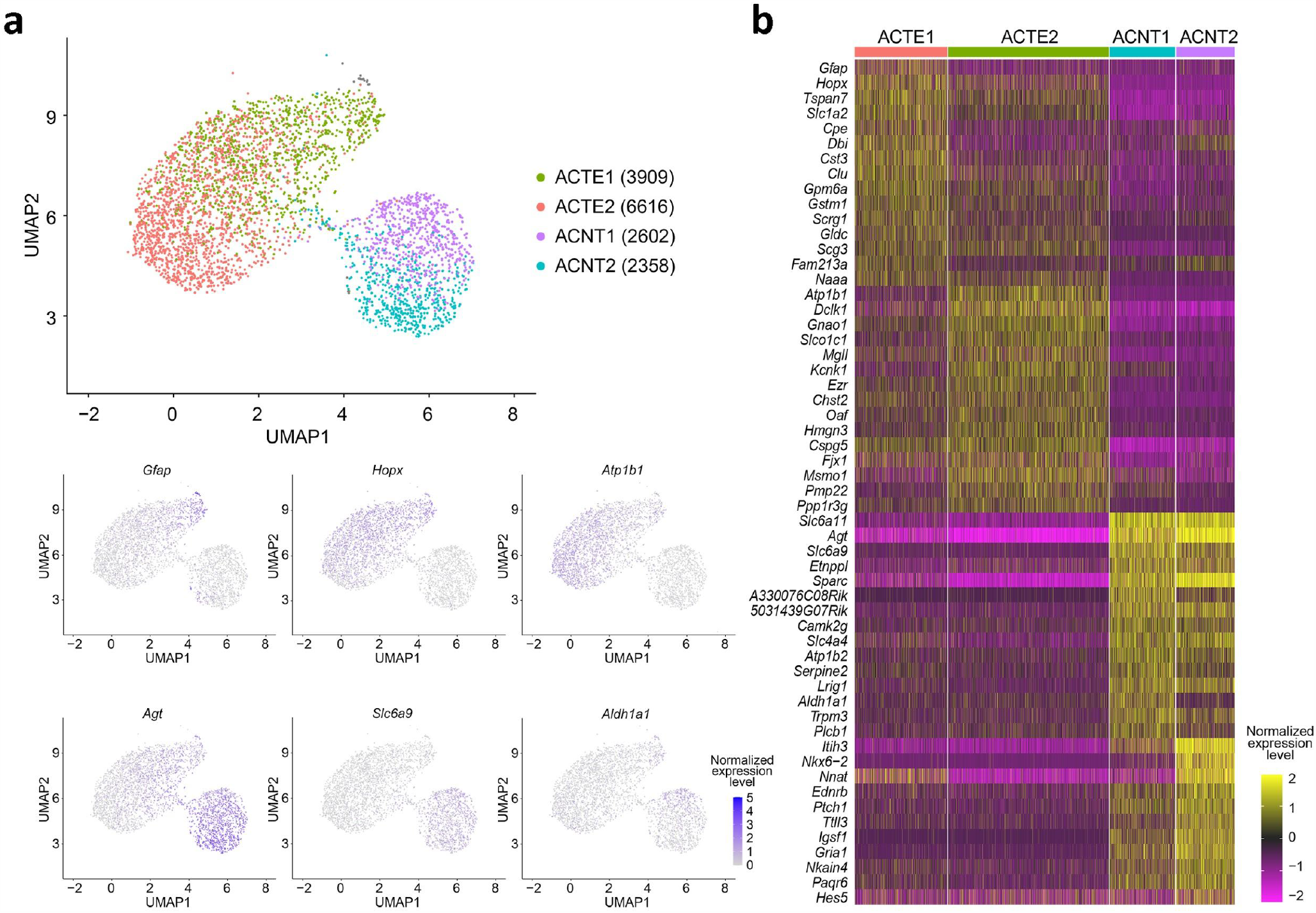
Identification of regional AC subtypes. **(a)** UMAP visualization and **(b)** heatmap of marker gene expression patterns in the different regional AC subtype clusters, including telencephalic AC cluster 1 (ACTE1), telencephalic AC cluster 2 (ACTE2), non-telencephalic AC cluster 1 (ACNT1) and non-telencephalic AC cluster 2 (ACNT2). Numbers in brackets: cell numbers for the respective AC subtypes. For clarity, 4000 cells were subsampled for visualization in each plot in **(a)**.

**Supplementary Fig. 3.**
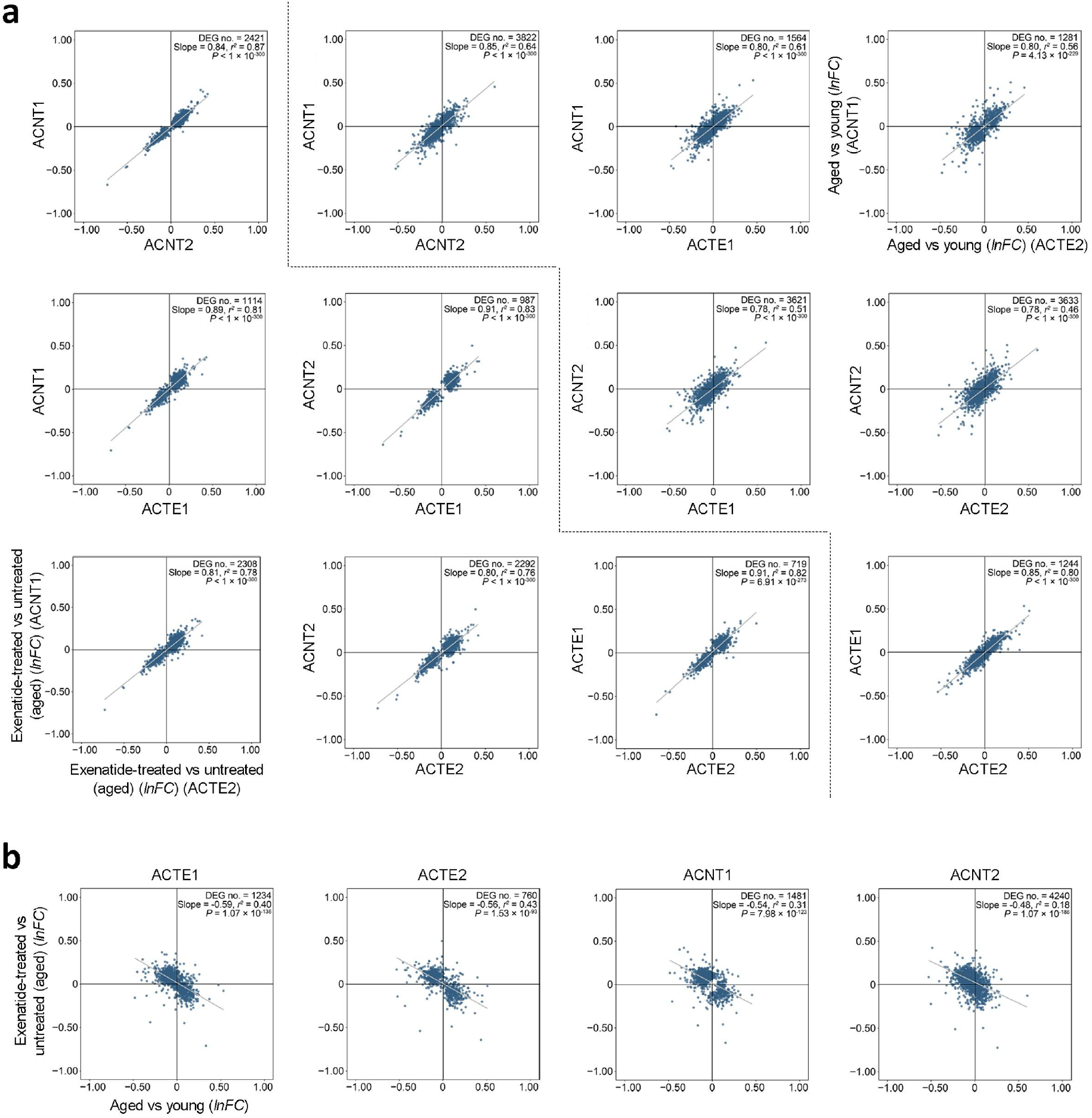
Consistency of differential expressions in the different regional AC subtype clusters in aging and after GLP-1RA treatment. **(a)** Pairwise comparisons of age-related (above-diagonal plots in the array) and GLP-1RA treatment-associated (below-diagonal plots in the array) differential expressions in the four AC subtypes (ACTE1, ACTE2, ACNT1 and ACNT2). **(b)** Expression changes in the regional AC subtypes in aging (*x*-axis) plotted against that after GLP-1RA treatment (*y*-axis). For all plots in **(a)** and **(b)**: Each dot represents one DEG. *lnFC*: natural log of fold change. Grey lines: lines of best fit by linear regression.

**Supplementary Fig. 4.**
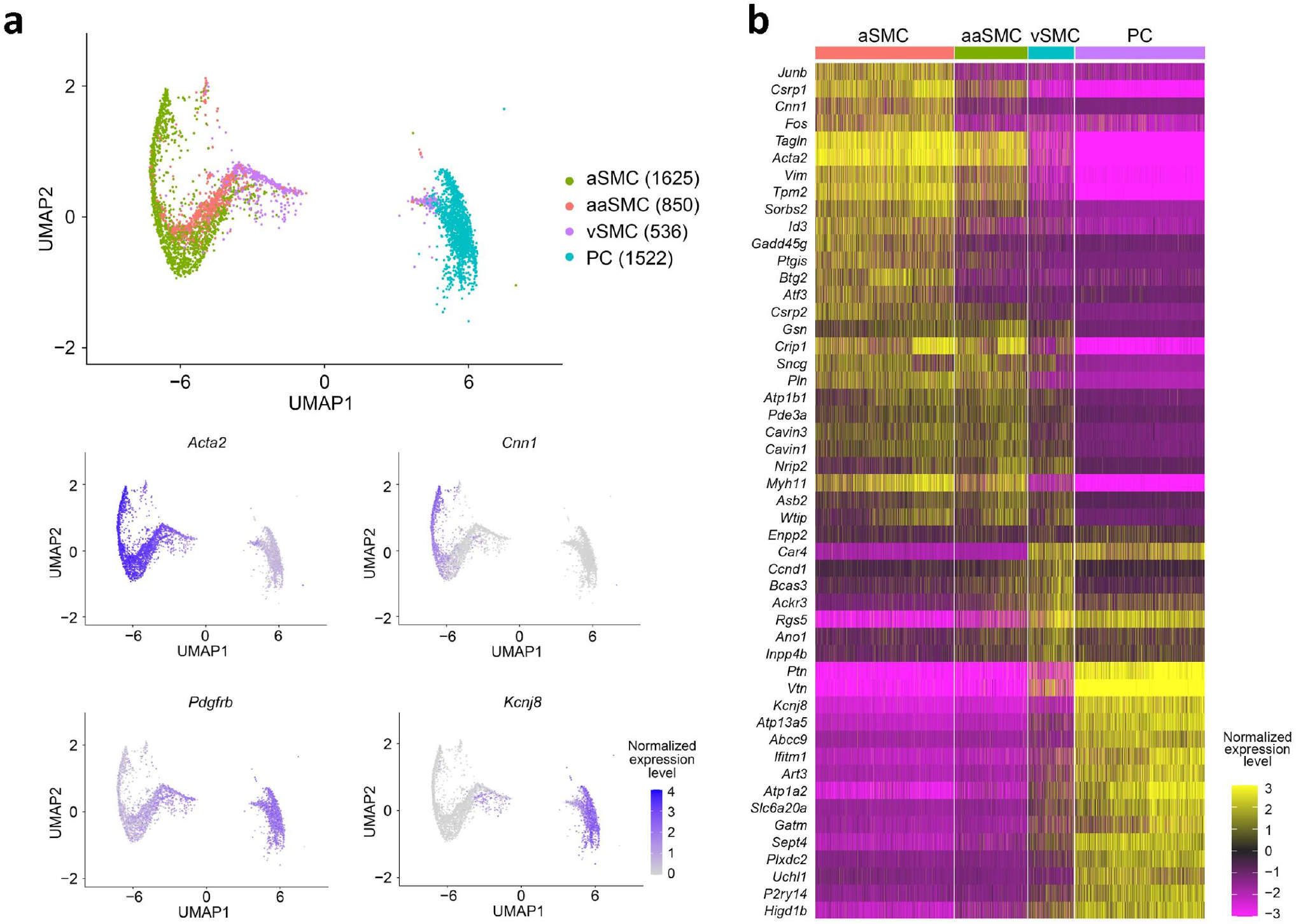
Identification of mural cell subtypes. **(a)** UMAP visualization and **(b)** heatmap of marker gene expression patterns in the different mural subtypes, including arterial SMC (aSMC), arteriolar SMC (aaSMC), venous SMC (vSMC) and pericyte (PC). Numbers in brackets: cell numbers for the respective cell subtypes.

**Supplementary Fig. 5.**
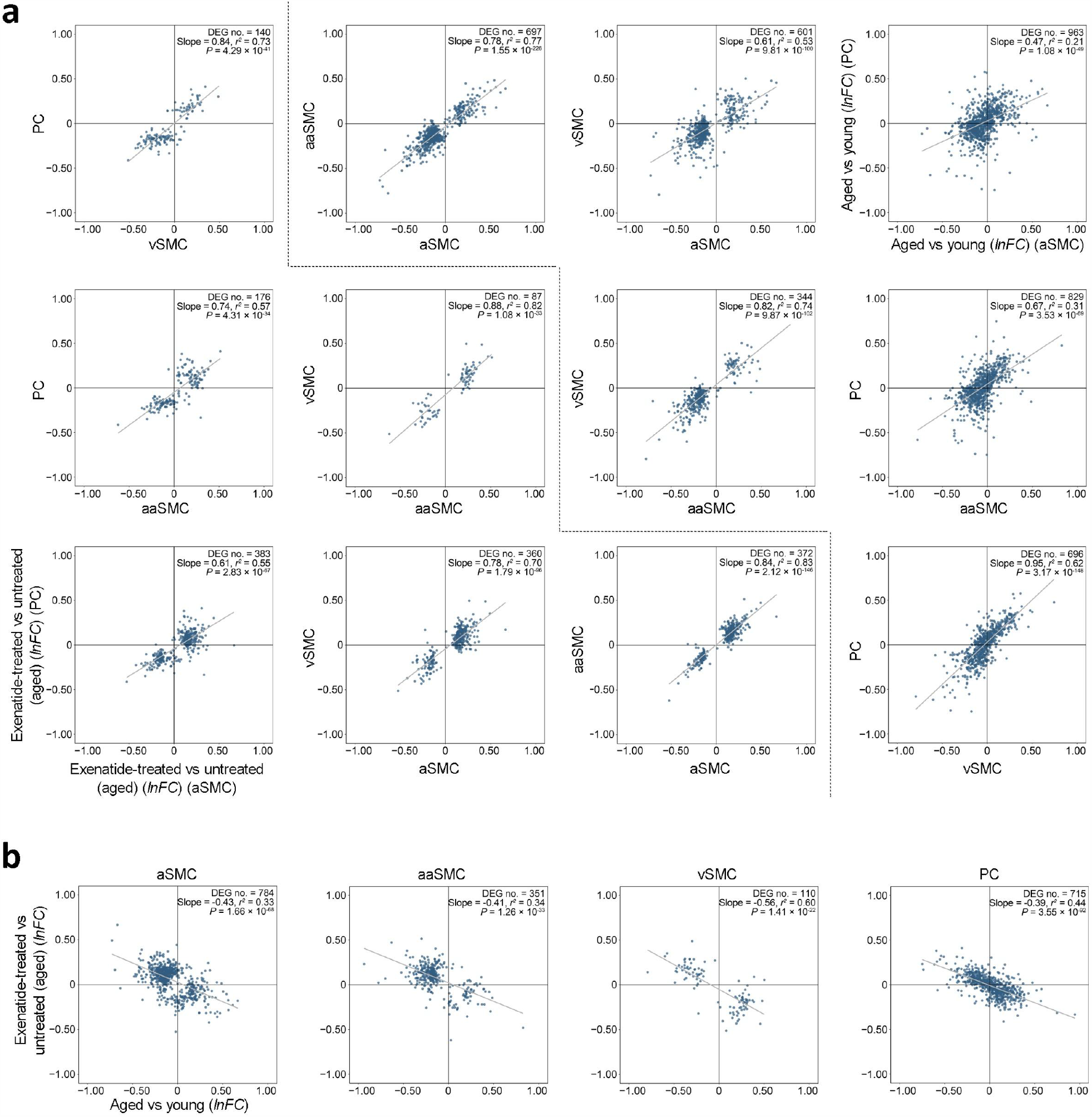
Consistency of differential expressions in the different mural cell subtypes in aging and after GLP-1RA treatment. **(a)** Pairwise comparisons of age-related (above-diagonal plots in the array) and GLP-1RA treatment-associated (below-diagonal plots in the array) differential expressions in the four mural cell subtypes (aSMC, aaSMC, vSMC and PC). **(b)** Expression changes in the mural cell subtypes in aging (*x*-axis) plotted against that after GLP-1RA treatment (*y*-axis). For all plots in **(a)** and **(b)**: Each dot represents one DEG. *lnFC*: natural log of fold change. Grey lines: lines of best fit by linear regression.

